# β-Blockers for ED Presentations with Recent Cocaine Use: A Systematic Review

**DOI:** 10.64898/2026.06.02.729530

**Authors:** Toby Paterson, Sai Vamshi Krishna Katraj, Aléchia Van Wyk, Vivetha Pooranachandran

## Abstract

**Background:** Cocaine produces cardiovascular toxicity through intense sympathetic stimulation and direct myocardial injury, generating presentations ranging from hypertension and tachycardia to coronary vasospasm, arrhythmia, and myocardial depression. β-blockers are foundational therapies in acute coronary syndromes, yet their use after cocaine exposure remains controversial due to concerns about unopposed α-adrenergic stimulation.

**Methods:** A systematic review was conducted in accordance with PRISMA guidelines. PubMed and Google Scholar (2000–2026) were searched for observational studies of adults (≥18 years) presenting to acute care with recent cocaine use that compared outcomes between β-blocker recipients and non-recipients. Eligible studies reported in-hospital mortality, myocardial infarction/troponin rise, clinically significant arrhythmia, or haemodynamic instability. Risk of bias was assessed using the Newcastle–Ottawa Scale, and certainty of evidence using GRADE.

**Results:** Four retrospective ED cohorts (n = 1,140) met inclusion criteria; 503 patients received at least one β-blocker dose. Across studies, β-blocker use was not associated with increased in-hospital mortality or malignant arrhythmias. Myocardial infarction was heterogeneous and sensitive to definition and timing. Haemodynamic data showed no hypertensive surge and modest systolic blood pressure reductions. Risk of bias was moderate, and certainty of evidence very low to low.

**Conclusions:** In typical ED presentations of recent cocaine use, β-blocker administration does not appear to increase mortality, myocardial infarction, or malignant arrhythmias, and available haemodynamic data do not support a reproducible unopposed-α response. However, mechanistic and preclinical evidence suggests potential harm in severe intoxication or myocardial depression. A selective, phenotype-guided approach is warranted, and prospective mechanistic studies are needed.

**Key Points:**

- Beta-blockers did not increase deaths, heart attacks, or dangerous heart rhythms in adults who came to the emergency department after recent cocaine use.
- Blood pressure generally decreased after beta-blocker treatment, and no consistent “unopposed alpha” reaction was seen in typical presentations.
- Caution is still needed in severe intoxication or cases with heart muscle weakness, but for most emergency presentations, beta-blockers appear safe when used appropriately.

## 1. Introduction

Cocaine-related presentations remain a frequent and clinically demanding problem in acute care. The drug produces a spectrum of cardiovascular complications, from tachyarrhythmias and hypertension to myocardial ischaemia and cardiac arrest, driven by intense sympathetic activation, coronary vasoconstriction and direct sodium-channel blockade [1]. These pathophysiological effects create a volatile clinical picture in which patients may present with isolated tachycardia, hypertensive crises, coronary vasospasm, malignant arrhythmia or, conversely, myocardial depression and shock [2,3]. Rising prevalence of cocaine use internationally, mean that cocaine-associated cardiovascular emergencies are an ongoing and growing burden for emergency clinicians [4].

Within this context, β-blockers occupy a uniquely controversial position. They are foundational therapies for angina, tachyarrhythmias and acute coronary syndromes, yet their use after cocaine exposure has been discouraged for decades because of the theoretical risk of unopposed α-stimulation [1-3]. That concern originated from early clinical reports and small experimental studies describing paradoxical hypertension and coronary vasoconstriction after propranolol in cocaine-exposed patients [4,5], and it has strongly influenced guidelines and clinical teaching despite limited mechanistic and clinical confirmation [6,7].

More recent observational work has challenged the universality of that caution. Several emergency-department cohorts and larger observational series have reported neutral or favourable outcomes after β-blocker administration in patients with cocaine-associated chest pain or recent use, with no consistent cardiovascular risk [6]. Nevertheless, the evidence remains fragmented: studies differ in how they define cocaine exposure (history or urine drug screen versus documented clinical intoxication), in outcome definitions (troponin rise versus adjudicated myocardial infarction), in timing of drug administration, and in the β-blocker classes evaluated [6].

This systematic review synthesises the clinical evidence most relevant to emergency practice and addresses a pragmatic question: in adults presenting to acute care with recent cocaine use, what does contemporary evidence demonstrate about the safety of β-blocker therapy? The review focuses on clinically meaningful outcomes, in-hospital mortality, myocardial infarction, arrhythmia, and haemodynamic instability, and highlights the methodological limitations that must inform interpretation and guide future research.

## 2. Methods

### 2.1 Protocols

This systematic review was conducted in accordance with the Preferred Reporting Items for Systematic Reviews and Meta-Analyses (PRISMA) guidelines [8].

### 2.2 Eligibility criteria

We included full-text, peer-reviewed observational studies of adults (≥18 years) presenting to acute care with recent cocaine use that compared clinical outcomes between patients who received a β-blocker during the index encounter (or at discharge) and those who did not. Eligible studies reported at least one of the following outcomes: in-hospital myocardial infarction (MI), all-cause mortality, clinically significant arrhythmia, and haemodynamic instability. We excluded studies that only reported outpatient or long-term β-blocker prescriptions without in-hospital exposure data.

### 2.3 Search Strategy

We searched PubMed, and Google Scholar from January 2000 to February 2026 using combined MeSH terms and keywords for cocaine, β-blockers (propranolol, metoprolol, labetalol, carvedilol, atenolol, esmolol), and acute toxicity/intoxication/overdose. (“Cocaine”[MeSH Terms] OR cocaine AND (“adrenergic beta antagonists”[MeSH Terms] OR betablocker *OR beta-blocker* OR propranolol OR labetalol OR metoprolol OR esmolol OR atenolol OR carvedilol AND, OR intoxication OR overdose OR poisoning OR acute.

Reference lists of relevant reviews and included articles were hand-searched for additional studies. Three reviewers (TP, SK and VP) independently screened titles and abstracts, retrieved full texts for potentially eligible reports, and resolved disagreements by discussion. A PRISMA flow diagram documents study selection.

### 2.4 Statistical Analysis

Quantitative pooling was not performed because the included studies differed substantially in exposure definition, β-blocker class, route, timing, and MI definition. These differences violate the assumptions required for meta-analysis, rendering heterogeneity statistics (I^2^), funnel plots, and Egger’s regression test inappropriate and potentially misleading. A narrative synthesis was therefore undertaken, in line with Cochrane and MOOSE guidance [8,9].

### 2.5 Risk of bias assessment

We assessed risk of bias for each observational study using the Newcastle–Ottawa Scale across selection, comparability, and outcome domains. Two reviewers (SK and VP) scored studies independently and resolved discrepancies by consensus. We considered confounding by indication, immortal-time bias (unclear timing of β-blocker relative to events), and outcome ascertainment (troponin-only vs clinically adjudicated MI) as key sources of bias and recorded these explicitly for each study.

### 2.6 Certainty of evidence

We assessed overall certainty for each outcome using GRADE principles adapted for observational evidence, considering risk of bias, inconsistency, indirectness (notably UDS vs documented intoxication), imprecision, and publication bias.

## 3. Results

Four observational ED/acute-care cohorts [10-13] met the inclusion criteria (Figure 1), comprising 1,140 adults in total. Across these cohorts 503 patients received β-blockers and 637 received alternative or no β-blocker therapy. Reported ages clustered around 40-50 years), and samples were predominantly male (Table 1). β-blocker exposure was heterogeneous but predominantly β1-selective agents (metoprolol); combined α/β agents (labetalol, carvedilol) were used in smaller proportions. Exact mg dosing was not reported and routes were mixed (IV and oral). Timing of administration was generally during the ED visit or early in admission, but precision of timing relative to biomarker sampling or events varied between studies (Table 2).

**Figure 1.**
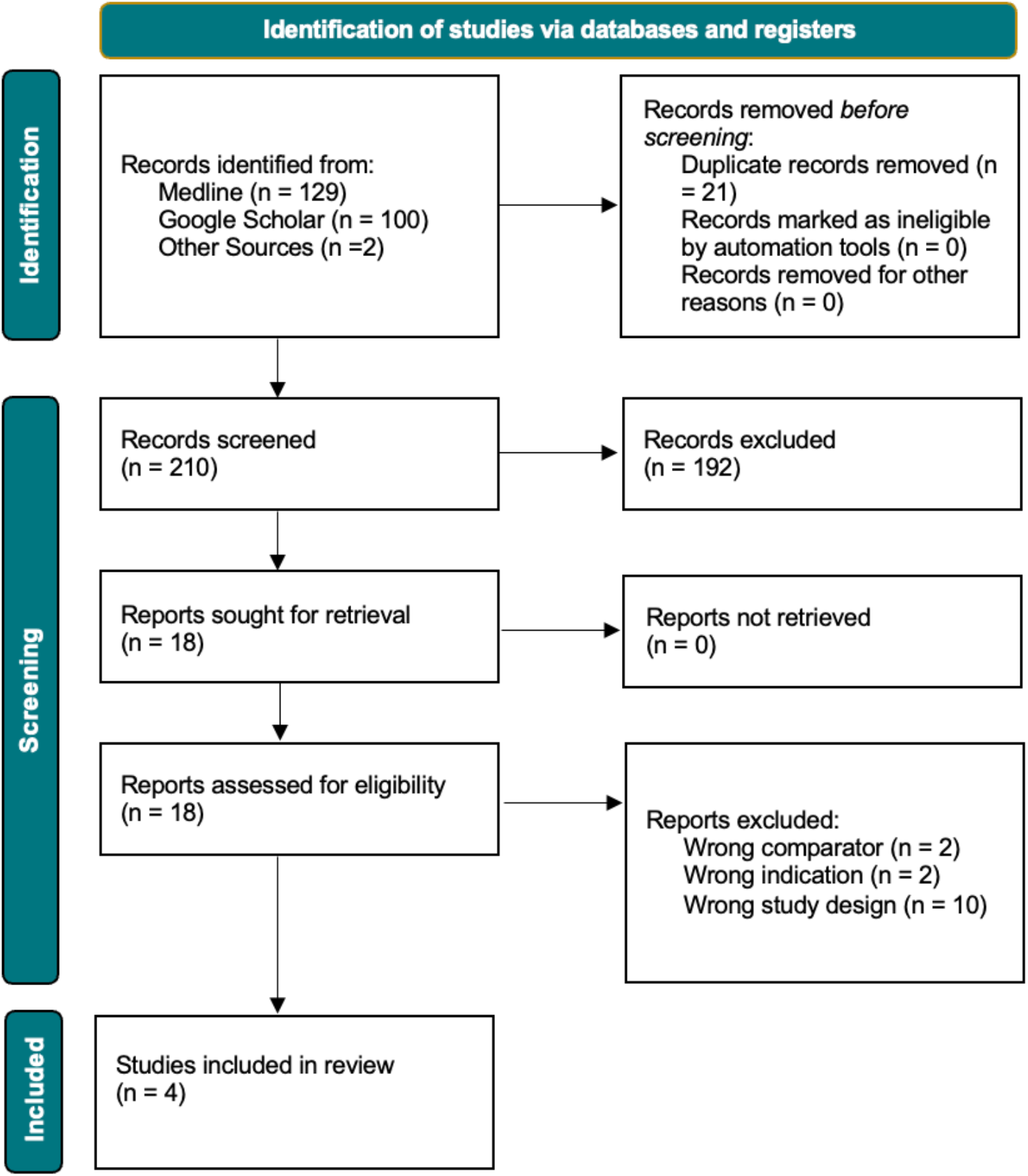
PRISMA flow diagram of study selection. Flow diagram illustrating the identification, screening, eligibility assessment, and inclusion of studies evaluating beta-blockers in the context of ED presentation with recent cocaine use. Following database searches and full-text review, 4 studies were included in the systematic review.

**Table 1:** Study characteristics for ED cohorts included in the systematic review [10-13]. ED: Emergency Department

| Study (year) | Country | Design | Population Size | Received β-blocker, n (%) | β-blocker | | No β-blocker | | Key aim / objective |
| --- | --- | --- | --- | --- | --- | --- | --- | --- | --- |
|  |  |  |  |  | Age | Male (%) | Age | Male (%) |  |
| **Rangel et al. (2010)** | USA | Retrospective Observational ED cohort | 328 | 151 (46) | 51 ± 9 | 111 (74) | 48 ± 9 | 124 (71) | Evaluate outcomes after β-blocker use in ED patients with recent cocaine use presenting with chest pain. |
| **Ibrahim et al. (2013)** | USA | Retrospective Observational ED cohort | 378 | 162 (43) | 46.2 ± 9 | 109 (67) | 42.3 ± 10.6 | 162 (75) | Assess troponin and clinical outcomes by β-blocker exposure and class in cocaine-positive ED patients. |
| **Fanari et al. (2014)** | USA | Retrospective Observational ED cohort | 376 | 164 (44) | 46 ± 9 | 112 (68) | 42 ± 11 | 157 (74) | Compare cardiovascular outcomes in cocaine-positive ED patients who did and did not receive β-blockers. |
| **Schmidt et al. (2015)** | USA | Retrospective Observational ED cohort | 58 | 26 (45) | 50 | 23 (89) | 48.3 | 30 (94) | Describe haemodynamic response and clinical outcomes after β-blocker administration in ED cocaine presentations. |

**Table 2:** β-blocker agents, dosing, administration route and key outcomes by study [10-13]. “β-blocker agent” denotes receipt of at least one β-blocker dose (IV or oral) during the index ED visit or subsequent admission. ED; Emergency Department, IV; Intravenous, PO; Oral administration (per os), n; number, IQR; Interquartile range, BP; Blood pressure, HR; Heart rate, MI; Myocardial infarction, VF; Ventricular fibrillation, VT; Ventricular tachycardia

| Study (year) | Positive Cocaine Test | β-blocker agent(s), n | Route (IV / PO), n | Timing to first dose (median, IQR) | Mortality, n | Myocardial infarction, n | Arrhythmia (ventricular / malignant events), n | Haemodynamic findings (BP/HR change) |
| --- | --- | --- | --- | --- | --- | --- | --- | --- |
| **Rangel et al. (2010)** | Metabolite benzoylcegonine in urine sample | IV Metoprolol = 113/151<br>Oral Metoprolol = 17/151<br>IV labetalol = 18/151 | IV = 131/151<br>PO = 22/151 | 100 minutes [49-279 minutes] in ED | β-blocker = 2/151<br>No β-blocker = 3/177<br>P = 0.76 | Troponin above 1.5ng/mL considered as acute MI.<br>β-blocker = 15/151 | VT or VF requiring electrical cardioversion or defibrillation.<br>β-blocker = 1/151 | Mean 8.6mmHg decrease in systolic BP (95% CI 14.7 to 2.5mm Hg) in β-blocker group (p = 0.006) |
| | | Oral labetalol = 3/151<br>Oral atenolol = 1/151<br>Oral propranolol = 1/151 | | 1070 minutes [691-1878 minutes] from triage to hospital ward | | No β-blocker = 18/177<br>P = 0.94 | No β-blocker = 1/177<br>P = 0.91 | |
| **Ibrahim et al. (2013)** | Metabolite benzoylecgonine in urine sample | Metoprolol = 88/162<br>Carvedilol = 44/162<br>Labetalol = 45/162<br>Atenolol = 4/162 | - | Within 24 hours | - | Myocardial necrosis defined as: Troponin I > 0.6ng/mL, and Troponin T > 0.1ng/mL<br>β-blocker = 22/162<br>No β-blocker = 20/216<br>P = 0.2 | - | No hypertensive emergencies reported (systolic BP >180mmHg; diastolic BP >110mmHg) |
| **Fanari et al. (2014)** | Urine toxicology screen | Metoprolol = 74/164<br>Carvedilol = 43/164<br>Labetalol = 44/164<br>Atenolol = 3/164 | - | Within 24 hours | β-blocker = 3/164<br>No β-blocker = 6/212<br>P = 0.13 | Troponin T > 0.1 ng/dL or significant ST elevation in 2 contiguous leads<br>β-blocker = 5/164<br>No β-blocker = 2/212<br>P = 0.18 | β-blocker = 7/164<br>No β-blocker = 5/212<br>P = 0.72 | - |
| **Schmidt et al. (2015)** | Urine test for metabolites | Oral Metoprolol = 13/26<br>IV Metoprolol = 5/26<br>IV Labetalol = 2/26<br>Oral Carvedilol = 2/26<br>Oral Atenolol = 2/26<br>Oral Propranolol = 2/26 | IV = 7/26<br>PO = 19/26 | During admission | β-blocker = 0/26<br>No β-blocker = 0/32 | Defined by positive Troponin I<br>β-blocker = 14/26<br>No β-blocker = 17/32 | - | Serial BP/HR: trend to lower BP/HR in β-blocker group over 6–12 h |

### 3.1 In-hospital mortality

Deaths were rare across studies. Reported in-hospital mortality counts for β-blocker vs no β-blocker were: Rangel et al. 2/151 vs 3/177; Fanari et al. 3/164 vs 6/212, and Schmidt et al. 0/26 vs 0/32. No study demonstrated a consistent increase in in-hospital all-cause mortality associated with β-blocker use; point estimates trend toward neutrality or fewer deaths with β-blockers, but confidence is limited by small event counts and retrospective designs.

### 3.2 Myocardial infarction

MI definitions varied substantially between studies, ranging from troponin-only criteria to combined clinical and ECG-based definitions. Across cohorts, there was no consistent signal of an increased risk of in-hospital MI attributable to β-blocker administration. However, results were heterogeneous and sensitive to both the MI definition used and the timing of β-blocker administration relative to biomarker sampling.

### 3.3 Arrhythmias

Arrhythmia reporting was sparse and lacked granularity. Across studies, there was no reproducible evidence of an increased incidence of malignant ventricular arrhythmias following β-blocker use. Interpretation remains limited by very small event numbers and non-adjudicated outcomes.

### 3.4 Haemodynamic responses and the unopposed-α hypothesis

Schmidt et al. observed falling BP and HR over 6–12 hours in β-blocker recipients, and Rangel et al. reported a mean early systolic BP reduction of ≈ −8.6 mmHg (95% CI −14.7 to −2.5). These findings do not support a clinically meaningful or reproducible unopposed-α hypertensive response in routine ED practice.

### 3.5 Risk of bias and certainty

Across the four included retrospective ED cohorts, NOS scores ranged from 5 to 6 out of 9, indicating moderate overall risk of bias (supplementary material, table 1). All studies were retrospective and at risk of confounding by indication, selection bias, and measurement heterogeneity. Key limitations: variable cocaine-exposure definitions (UDS vs clinical intoxication), inconsistent MI definitions (troponin vs clinical/ECG), incomplete reporting of BB dose and exact timing, and sparse arrhythmia adjudication. Overall certainty of evidence is low to very low (supplementary material, table 2).

Across the four included observational cohorts, β-blocker administration (mostly metoprolol; some labetalol/carvedilol) was not associated with a reproducible increase in in-hospital mortality or malignant arrhythmias, and the best haemodynamic data did not show the expected hypertensive “unopposed-α” surge. Myocardial infarction/troponin results were heterogeneous and therefore inconclusive. Given retrospective designs, small event counts, and variable definitions, these findings are hypothesis-generating and should be interpreted with caution.

## 4. Discussion

Cocaine cardiotoxicity arises from the convergence of two dominant mechanisms: a systemic catecholamine surge driven by monoamine reuptake inhibition, producing α-mediated vasoconstriction and β-mediated tachycardia and inotropy, and direct myocardial cellular injury resulting from sodium-channel blockade, impaired sarcoplasmic-reticulum calcium handling and local anaesthetic effects on depolarisation [1-3]. The clinical phenotype at any moment, hypertension, tachycardia, coronary vasospasm, ventricular arrhythmia or myocardial depression, reflects the shifting balance between these processes and the patient’s physiological reserve [14-16]. This dual-pathway model explains why β-blockade may be beneficial when sympathetic overdrive predominates yet potentially harmful when direct myocardial toxicity is the dominant driver [1].

There is a mechanistic rationale for β-blocker therapy in patients whose presentation is characterised by adrenergic excess. β_1_-selective blockade reduces heart rate, contractility and myocardial oxygen demand, and dampens β-adrenergic proarrhythmic signalling [1,2]. Combined α/β-blockers such as labetalol additionally counteract α_1_-mediated vasoconstriction and coronary spasm, offering a more comprehensive approach in patients with hypertension and tachycardia [17,18]. These mechanisms align with clinical observations from case reports describing rapid haemodynamic improvement after intravenous labetalol or metoprolol in topical cocaine toxicity and stimulant co-ingestion, with marked reductions in systolic blood pressure and heart rate [17,19].

However, preclinical data delineate a boundary beyond which β-blockade may be deleterious. In a chronically instrumented LD_50_ rat model, administration of propranolol or labetalol after rapid intravenous cocaine exposure worsened survival, suggesting that adrenergic blockade can remove essential compensatory support during overwhelming catecholamine storms [20]. Isolated myocardial tissue experiments similarly demonstrate that cocaine-induced negative inotropy is amplified by β_1_-blockade and reversible with extracellular calcium, implicating impaired calcium release and sodium-channel effects [1]. Central monoaminergic modulation further complicates prediction of haemodynamic responses, as cocaine can indirectly activate inhibitory α_2_ and 5-HT_1_A pathways, meaning the net effect of β-blockade depends on the prevailing balance of central and peripheral receptor activity [21].

Clinical cases illustrate these mechanistic boundaries. A recent report of carvedilol–colchicine–cocaine co-ingestion described an initial period in which cocaine’s adrenergic stimulation masked β-blocker toxicity, followed by delayed bradycardia, hypotension, cardiogenic pulmonary oedema and multiorgan failure [22]. The combination of direct myocardial depression, co-ingestant toxicity and delayed β-blocker effects produced a clinical trajectory fundamentally different from the presentations typically seen in emergency-department (ED) cohorts. Such cases highlight that β-blockade may be hazardous in severe intoxication, large overdose or presentations dominated by myocardial depression rather than sympathetic excess [23].

ED cohort data must therefore be interpreted within this mechanistic context. The four included studies [10-13], enrolled patients with cocaine-associated chest pain and recent use identified by history or urine drug screen, rather than individuals in a fulminant catecholamine storm. Across these cohorts, β-blocker exposure, most often with β_1_-selective agents, was not associated with hypertensive rebound, increased in-hospital mortality or excess malignant arrhythmias. Serial haemodynamic measurements in Schmidt et al.’s study showed no evidence of unopposed α-stimulation, and Rangel et al. reported modest reductions in systolic blood pressure after early β-blocker use [10,13]. Arrhythmic outcomes were similarly neutral, with no reproducible signal of harm [10,12]. These findings show that in heterogeneous, lower-severity ED populations, where sympathetic activation predominates, β-blockade more commonly attenuates adrenergic stress than unmasks direct myocardial depression.

Meta-analytic data reinforce this interpretation. A 2019 systematic review by Shin and colleagues, including eight observational studies and over 2,000 patients, found no significant difference in in-hospital mortality (RR 0.59, 95% CI 0.24–1.47) or myocardial infarction (RR 1.24, 95% CI 0.74–2.06) between β-blocker users and non-users [6]. Long-term outcomes were similarly neutral. A separate meta-analysis by Pham et al. and Lo et al. reported no increase in non-fatal myocardial infarction (OR 1.36, 95% CI 0.68–2.75; RR 1.08, 95% CI 0.61-1.91) or all-cause mortality (OR 0.68, 95% CI 0.26–1.79; RR 0.75, 95% CI 0.46-1.24), respectively [24,25]. These pooled findings are consistent with the absence of harm in ED cohorts, although heterogeneity in myocardial infarction definitions and timing of β-blocker administration limits precision.

Older studies contribute important nuance. Mohamad et al. reported a higher rate of myocardial necrosis in β-blocker recipients, whereas Dattilo et al. found a markedly lower risk of myocardial infarction associated with β-blocker use [26,27]. These contradictory findings likely reflect confounding by indication, differences in troponin thresholds and variation in β-blocker class. Early haemodynamic studies also shaped guideline caution. Lange and colleagues demonstrated potentiation of cocaine-induced coronary vasoconstriction with intracoronary propranolol, whereas Boehrer et al. found that intravenous labetalol attenuated vasoconstriction [5,28]. These studies used small samples, non-selective agents and routes of administration not reflective of clinical practice, yet they continue to influence perceptions of risk.

Agent selection and clinical severity remain decisive modifiers of net effect. Combined α/β-blockers are mechanistically preferable when hypertension and tachycardia coexist, whereas β_1_-selective agents are appropriate for isolated rate control in haemodynamically stable patients [28]. Non-selective β_1_/β_2_ blockers carry the greatest theoretical risk of removing β_2_-mediated vasodilation and have the strongest animal signals of harm [20,29,30]. Regardless of agent, cautious titration with continuous ECG and blood-pressure monitoring is prudent so that any unanticipated negative inotropic response can be detected and managed.

The current evidence base limits definitive mechanistic conclusions. Human studies are predominantly retrospective and vulnerable to confounding by indication; many define exposure by urine drug screen rather than contemporaneous intoxication, and haemodynamic and arrhythmic endpoints are inconsistently reported [10-13]. Animal and tissue models use supraphysiological exposures and lack human compensatory mechanisms; they illuminate plausible mechanisms of catastrophic harm but do not determine typical clinical outcomes [1,20, 29, 30]. Case reports highlight rare but severe presentations that are not represented in ED cohorts [22]. These limitations underscore the need for caution in extrapolating cohort safety to patients with severe intoxication, seizures, respiratory compromise or large overdose.

Resolving remaining uncertainty will require prospective mechanistic human studies. Priority designs include enrolment of patients with documented clinical intoxication, continuous haemodynamic and ECG monitoring, and predefined agent and dose protocols. Comparative studies of β_1_-selective versus combined α/β agents, stratified by severity would clarify class-specific effects. Translational work examining cocaine’s impact on sarcoplasmic-reticulum calcium handling and stereoisomer-specific β-blocker pharmacology may further refine mechanistic understanding. Until such data are available, β-blocker therapy should be considered selectively, guided by clinical phenotype, haemodynamic stability and awareness of the mechanistic boundaries that distinguish typical ED presentations from severe intoxication.

## 5. Conclusion

This review found no clear evidence of increased harm from β-blocker use in typical ED presentations of cocaine exposure, nor convincing evidence of overall benefit; preclinical models, however, delineate a plausible high-risk boundary in which β-blockade may be deleterious. Taken together, the data support selective caution rather than categorical prohibition: favour β1-selective or combined α/β agents according to the clinical phenotype, titrate carefully under continuous monitoring, and avoid routine use of nonselective β1/β2 agents in patients with suspected overdose, seizures, or respiratory compromise. Definitive guidance will require prospective, mechanistic human studies with documented intoxication, continuous haemodynamic monitoring, and randomised comparisons of agent classes.

