## Supplementary Material for "β-Blockers for ED Presentations with Recent Cocaine Use: A Systematic Review"

| **Study** | **Selection (0–4)** | **Comparability (0–2)** | **Outcome (0–3)** | **Total (0–9)** | **Key Limitations** |
| --- | --- | --- | --- | --- | --- |
| **Rangel et al. (2010)** | 3 | 1 | 2 | **6** | Confounding by indication; UDS‑only exposure; unclear β‑blocker timing → immortal‑time bias; MI defined by troponin only; sparse arrhythmia adjudication |
| **Ibrahim et al. (2013)** | 3 | 1 | 2 | **6** | Troponin‑only MI definition; incomplete timing of β‑blocker; UDS‑based exposure; limited adjustment for confounders; no adjudicated arrhythmia outcomes |
| **Fanari et al. (2014)** | 3 | 1 | 2 | **6** | UDS‑based exposure; mixed β‑blocker classes; MI definition inconsistent; β‑blocker timing “within 24h” → risk of immortal‑time bias; limited arrhythmia adjudication |
| **Schmidt et al. (2015)** | 2 | 1 | 2 | **5** | Very small sample; UDS‑only exposure; unclear timing; no mortality/arrhythmia adjudication; troponin‑only MI |

**Table 1. Newcastle–Ottawa Scale (NOS) assessment of included ED cohort studies [10-13].**

The NOS evaluates observational study quality across three domains: Selection (4 items), Comparability (2 items), and Outcome (3 items). Each item is awarded one point if criteria are met, with a maximum score of 9. Higher scores indicate lower risk of bias. Key sources of bias relevant to this review include confounding by indication, variable cocaine exposure definitions, inconsistent myocardial infarction criteria, unclear β‑blocker timing leading to potential immortal‑time bias, and non‑adjudicated arrhythmia outcomes. Total scores reflect the overall methodological robustness of each study. UDS; Urine drug screen, MI; Myocardial infarction

| **Outcome** | **Studies contributing** | **Effect estimate / direction** | **Key design features** | **Main risks of bias (with concrete details)** | **Indirectness (what is indirect)** | **Imprecision (numbers & CIs)** | **Overall GRADE rating** |
| --- | --- | --- | --- | --- | --- | --- | --- |
| **All‑cause in‑hospital mortality** | [10-13] | No clear difference between β‑blocker and no β‑blocker; absolute event rates very low in all cohorts | All 4 are retrospective ED cohorts of UDS‑positive cocaine chest pain; β‑blocker exposure not randomised; treatment at clinician discretion | **Confounding by indication:** β‑blocker patients older, more HTN, more CAD, higher TIMI risk, more intensive care [10-12].    **Immortal‑time bias:** β‑blockers often given hours after triage (median 100 min in ED; 1070 min to ward dose) [10].  **Outcome ascertainment:** mortality captured only in‑hospital; no blinded adjudication; small numbers of deaths (e.g. 3 vs 6 [12]; 2 vs 3 [10]). | **Population:** UDS‑positive, not necessarily acutely intoxicated; timing of cocaine use often self‑reported or unknown.  **Intervention:** mix of β1‑selective and α/β agents; dose and exact timing variably reported.  **Outcome:** short‑term in‑hospital mortality only; no long‑term follow‑up in [11-13]. | Very few deaths per study; CIs were wide and compatible with both harm or benefit (e.g. [12] | **VERY LOW** |
| **Myocardial infarction / myocardial necrosis** | [10-13] | No consistent difference; some numerically higher MI in β‑blocker group, but non‑significant after adjustment | Same 4 retrospective ED cohorts | **Confounding by indication:** higher baseline risk in β‑blocker group (more CAD, LV dysfunction, higher BP, more typical angina).  **Outcome definition heterogeneity:** MI = troponin I >1.5 ng/mL [10]; MI = troponin T >0.1 ng/dL or ST‑elevation [12]; “myocardial necrosis” = troponin I >0.6mg/dL or T >0.1mg/dL [11]; [13] reports “positive troponin” without universal MI criteria.  **No blinded adjudication**: timing of β‑blocker vs troponin rise unclear in most. | **Population:** UDS‑positive chest pain, not necessarily acute intoxication; some chronic cocaine users.  **Outcome:** biochemical MI surrogates rather than clinically adjudicated type 1 MI; no systematic ECG/imaging confirmation.  **Intervention:** mixed β‑blocker classes; no stratified effect by acute vs remote cocaine use. | Event counts modest (e.g. MI: 5 vs 2 [12]; 22 vs 20 with troponin rise [11]; 15 vs 18 [10]; CIs wide (e.g. Composite OR 1.37, 95% CI 0.64–2.93 [12]; Underpowered to exclude clinically important harm [11]. | **VERY LOW** |
| **Ventricular arrhythmia / malignant events** | [10,12,13] | Very rare events; no clear difference between groups | Retrospective ED cohorts; arrhythmias captured from telemetry/clinical documentation | **Detection bias:** arrhythmias identified from routine clinical care, not systematic monitoring; no central adjudication.  **Confounding by indication:** higher‑risk patients more likely to receive β‑blockers and more intensive monitoring.  **Sparse data:** e.g. VT/VF 1 vs 1 [10]; 7 vs 5 [12]; No detailed arrhythmia breakdown [13]. | **Population:** UDS‑positive chest pain; arrhythmia risk may differ in acutely intoxicated vs remote users.  **Outcome:** only in‑hospital events; no out‑of‑hospital or post‑discharge arrhythmias; definitions limited to VT/VF requiring intervention. | Very small number of events across all studies; no effect estimate with narrow CI; any pooled estimate would have extremely wide CI and be unstable. | **VERY LOW** |
| **Haemodynamic response (BP/HR change)** | [10,12,13] | β‑blockers associated with modest reductions in BP and HR; no signal of “unopposed α” hypertension | Same retrospective cohorts; haemodynamic data derived from routine vitals | **Confounding by indication:** β‑blocker patients more hypertensive and tachycardic at baseline; other antihypertensives co‑administered (nitrates, ACEI, benzodiazepines).  **Timing:** vitals often measured after multiple interventions; not isolated β‑blocker effect. | **Population:** ED chest pain with positive UDS; haemodynamic response may differ in pure intoxication vs mixed pathology (HTN, LV dysfunction).  **Outcome:** short‑term BP/HR changes only; no direct patient‑important endpoint. | Adjusted mean systolic BP change −8.6 mmHg (95% CI −14.7 to −2.5) favouring ED β‑blocker [10]; Similar or lower BP/HR over 6–12 h in β‑blocker group; sample sizes modest but CIs reasonably tight for BP change [13]. | **LOW** |

**Table 2. GRADE evidence profile for β‑blocker use in cocaine‑associated chest pain [10-13].**

Certainty of evidence was assessed using the GRADE framework across five domains: risk of bias, inconsistency, indirectness, imprecision, and publication bias. Evidence from observational studies begins at *low certainty* and is downgraded where concerns are identified. Risk of bias was downgraded for all clinical outcomes due to retrospective design, confounding by indication, UDS‑based exposure misclassification, inconsistent MI definitions, and absence of outcome adjudication. Indirectness reflects uncertainty regarding acute intoxication status and variability in β‑blocker timing. Imprecision reflects small event numbers and wide confidence intervals. Overall certainty was rated very low for mortality, myocardial infarction, and ventricular arrhythmia, and low for haemodynamic outcomes.

GRADE; Grading of Recommendations, Assessment, Development and Evaluation, ED; Emergency Department, UDS; Urine drug screen, MI; Myocardial infarction, CAD; Coronary artery disease, HTN; Hypertension, VF; Ventricular fibrillation, VT; Ventricular tachycardia, TIMI; Thrombolysis in Myocardial Infarction (risk score), BP; Blood pressure, HR; Heart rate, ACEI; Angiotensin‑converting enzyme inhibitor, LV; Left Ventricular, CI; Confidence interval, OR; Odds ratio.
